# Surface colonization by *Flavobacterium johnsoniae* promotes its survival in a model microbial community

**DOI:** 10.1101/2024.01.05.574367

**Authors:** Shruthi Magesh, Amanda I. Hurley, Julia F. Nepper, Marc G. Chevrette, Jonathan H. Schrope, Chao Li, David J. Beebe, Jo Handelsman

## Abstract

*Flavobacterium johnsoniae* is a ubiquitous soil and rhizosphere bacterium, but despite its abundance, the factors contributing to its success in communities are poorly understood. Using a model microbial community, <u>T</u>he <u>H</u>itchhikers <u>O</u>f the Rhizosphere (THOR), we determined the effects of colonization on fitness of *F. johnsoniae* in the community. Insertion sequencing (INSeq), a massively parallel transposon mutant screen, on sterile sand identified 25 genes likely to be important for surface colonization. We constructed in-frame deletions of nine candidate genes predicted to be involved in cell membrane biogenesis, motility, signal transduction, and transport of amino acids and lipids. All mutants poorly colonized sand, glass, and polystyrene and produced less biofilm than the wild type, indicating the importance of the targeted genes in surface colonization. Eight of the nine colonization-defective mutants were also unable to form motile biofilms, or zorbs, thereby suggesting that the affected genes play a role in group movement and linking stationary and motile biofilm formation genetically. Furthermore, we showed that deletion of colonization genes in *F. johnsoniae* affected its behavior and survival in THOR on surfaces, suggesting that the same traits are required for success in a multispecies microbial community. Our results provide insight into the mechanisms of surface colonization by *F. johnsoniae* and form the basis for further understanding its ecology in the rhizosphere.

**IMPORTANCE:** Microbial communities direct key environmental processes through multispecies interactions. Understanding these interactions is vital for manipulating microbiomes to promote health in human, environmental, and agricultural systems. However, microbiome complexity can hinder our understanding of the underlying mechanisms in microbial community interactions. As a first step towards unraveling these interactions, we explored the role of surface colonization in microbial community interactions using THOR, a genetically tractable model community of three bacterial species, *Flavobacterium johnsoniae*, *Pseudomonas koreensis,* and *Bacillus cereus.* We identified *F. johnsoniae* genes important for surface colonization in solitary conditions and in the THOR community. Understanding the mechanisms that promote success of bacteria in microbial communities brings us closer to targeted manipulations to achieve outcomes that benefit agriculture, the environment, and human health.

## INTRODUCTION

Microorganisms are essential in every ecosystem and often exist in communities where they interact with each other and the environment (1). The soil microbiome drives key biogeochemical cycles, and the rhizosphere microbiome influences plant susceptibility to disease and drought, indicating the critical role of microbial communities in environmental health and agricultural productivity (2, 3). The rhizosphere is the region in soil that is influenced by nutrient-rich plant root secretions, or root exudate, which attract microorganisms from the surrounding soil (3). Rhizosphere microorganisms are essential to plant health as they promote plant nutrient uptake, stress tolerance, and disease suppression (4), and yet we know relatively little of the genes involved in mediating rhizosphere colonization by many soil bacteria. Hence, understanding microbial colonization in this dynamic environment is an important step toward modifying the soil microbiome for agricultural benefit.

After decades of studying bacterial monocultures, the field of microbiology has begun to explore the significance of multispecies interactions. Advances in high-throughput sequencing technology have enabled the use of metagenomics and metatranscriptomics to determine the taxonomic composition and functional characteristics of microbiomes in soil and rhizospheres (5–9). Nevertheless, the complexity of microbiomes containing thousands of interacting species makes it difficult to determine functions of genes using classical genetic approaches.

Genetic analysis of a simplified model microbial community enables mechanistic understanding of community interactions. We previously described <u>T</u>he <u>H</u>itchhikers <u>O</u>f the <u>R</u>hizosphere (THOR), a three-species model community composed of *Pseudomonas koreensis*, *Bacillus cereus*, and *Flavobacterium johnsoniae* isolated from the rhizospheres of field-grown alfalfa and soybean plants (10). These three species are genetically tractable, making mutant analysis in a community context possible. Additionally, numerous interactions between THOR members have been observed in both field and laboratory settings. For example, *P. koreensis* and *F. johnsoniae* were physically associated as biological hitchhikers with *B. cereus* isolated from soybean roots (10). The three species form more biofilm together than any of the species alone or in pairs (10). *B. cereus* provides peptidoglycan fragments as a carbon source that enables *F. johnsoniae* to grow in soybean root exudate (11). *F. johnsoniae* and *P. koreensis* induce *B. cereus* colony expansion (10). *F. johnsoniae* growth is inhibited by koreenceine, an antibiotic produced by *P. koreensis*, and koreenceine levels are modulated by the third THOR member, *B. cereus* (12). Gene expression and metabolite profiles of each member differ when they are in the community versus in pure culture (13, 14). These interactions indicate that THOR members engage in a web of biological interactions.

*Flavobacterium* spp. are abundant in the rhizosphere and are well studied for their ability to glide on surfaces using cell-surface motility adhesins that are delivered by a unique Type 9 Secretion System (T9SS) (15–18). Some strains of *F. johnsoniae* are even known to promote plant growth and suppress pathogens in soil but its behavior in the rhizosphere or on soil particles is poorly understood (19, 20). The elegant work of the McBride lab has provided the framework for genetic analysis in *F. johnsoniae* (21), enabling us to pinpoint genes important for surface colonization, which emerged as a critical factor in *F. johnsoniae* survival in the microbial community and biofilm formation on surfaces.

## RESULTS

### Genetic determinants of surface colonization in *F. johnsoniae*

We hypothesized that surface colonization promotes *F. johnsoniae* success in THOR. Here, we define surface colonization as the ability of bacteria to attach, proliferate, and survive on a substrate. To study surface colonization, we used sand, an abundant, easily manipulated soil particle that *Flavobacterium* spp. colonize better than other soil particles (22). We leveraged the power of insertion sequencing (INSeq) and conducted a massively parallel screen of *F. johnsoniae* transposon mutants to identify genes important for sand colonization. We created a library of approximately 75,000 transposon mutants using a modified *Bacteroides thetaiotaomicron* vector (pSAMFjoh 2) and compared mutant frequencies after growth under planktonic conditions (“input population”) with their frequencies after growth on sand (“output population”) for 48 h. By comparing the abundance of each transposon insertion between the planktonic and sand-colonized populations, we identified 25 candidate genes important for sand colonization. Mutants with transposon insertions in 21 different genes were underrepresented with a minimum fold change (planktonic/sand) of log2 > 1 and adjusted p-value < 0.05, suggesting that deletion of these genes would reduce sand colonization. Mutants with transposon insertions in four genes were overrepresented with a minimum fold change (planktonic/sand) of log2 < -1 with an adjusted p-value < 0.05, suggesting that deletion of these genes would increase colonization (Fig. 1A, Table 1 and Table S1).

**Figure 1:**
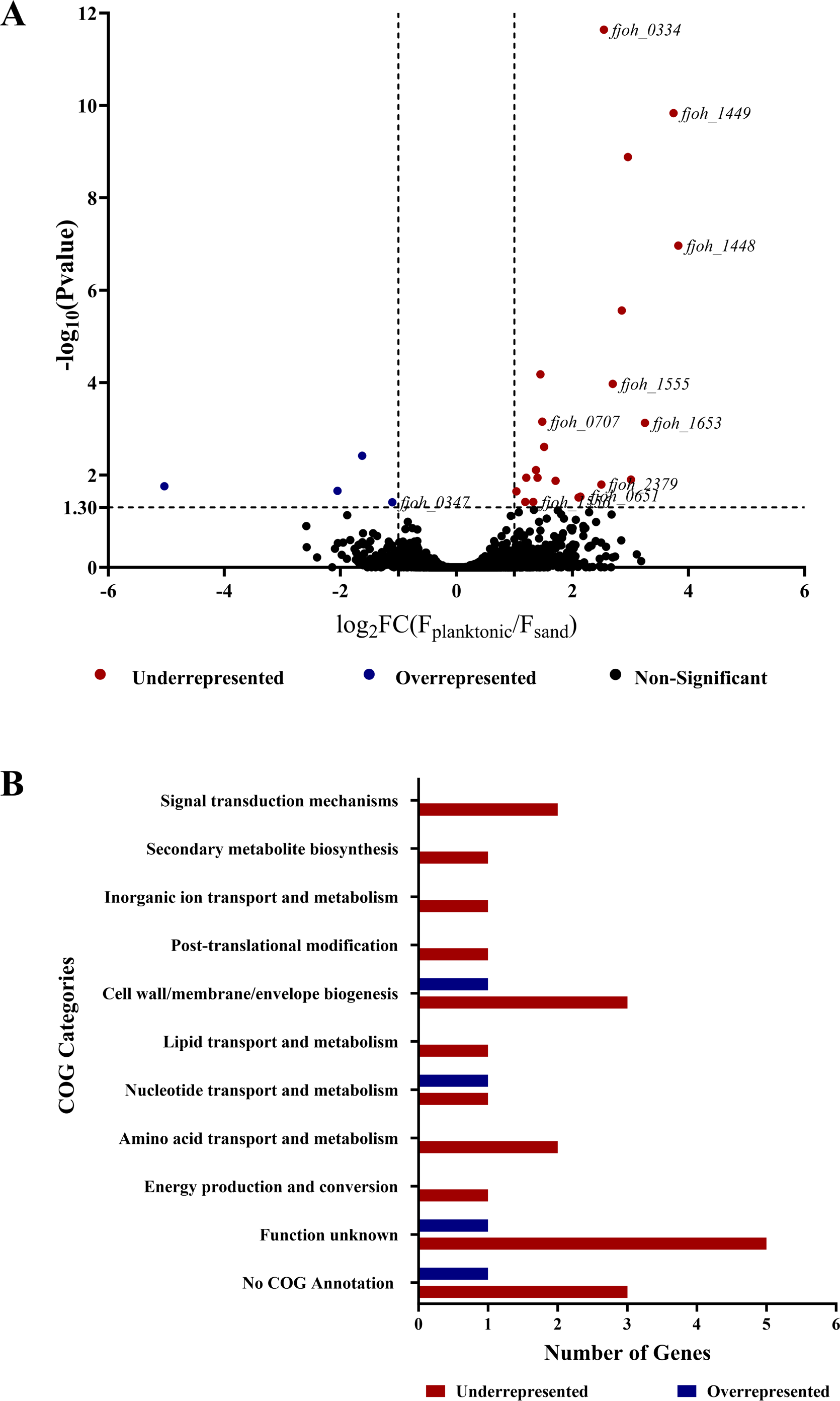
Genes identified by INSeq screen as important for sand colonization. (A) The relative abundance of transposon insertions for each gene was compared between *F. johnsoniae* grown planktonically and on sand. The data are presented as a volcano plot with –log_10_ p-value on the y-axis and log_2_ fold-change (input/output) on the x-axis. The black dots represent mutants whose representation did not differ between conditions (p-value >0.05), the red represent mutants that are significantly underrepresented on the surface of sand and the blue represent mutants that are significantly overrepresented on the surface of sand (p-value <0.05 and FC>1 or FC<-1). The genes affected in mutants whose representation differed between the two conditions were validated by gene deletions; these mutants are labeled by their Gene ID. (B) The genes identified as important in sand colonization are categorized based on COG annotation.

**Table 1:**
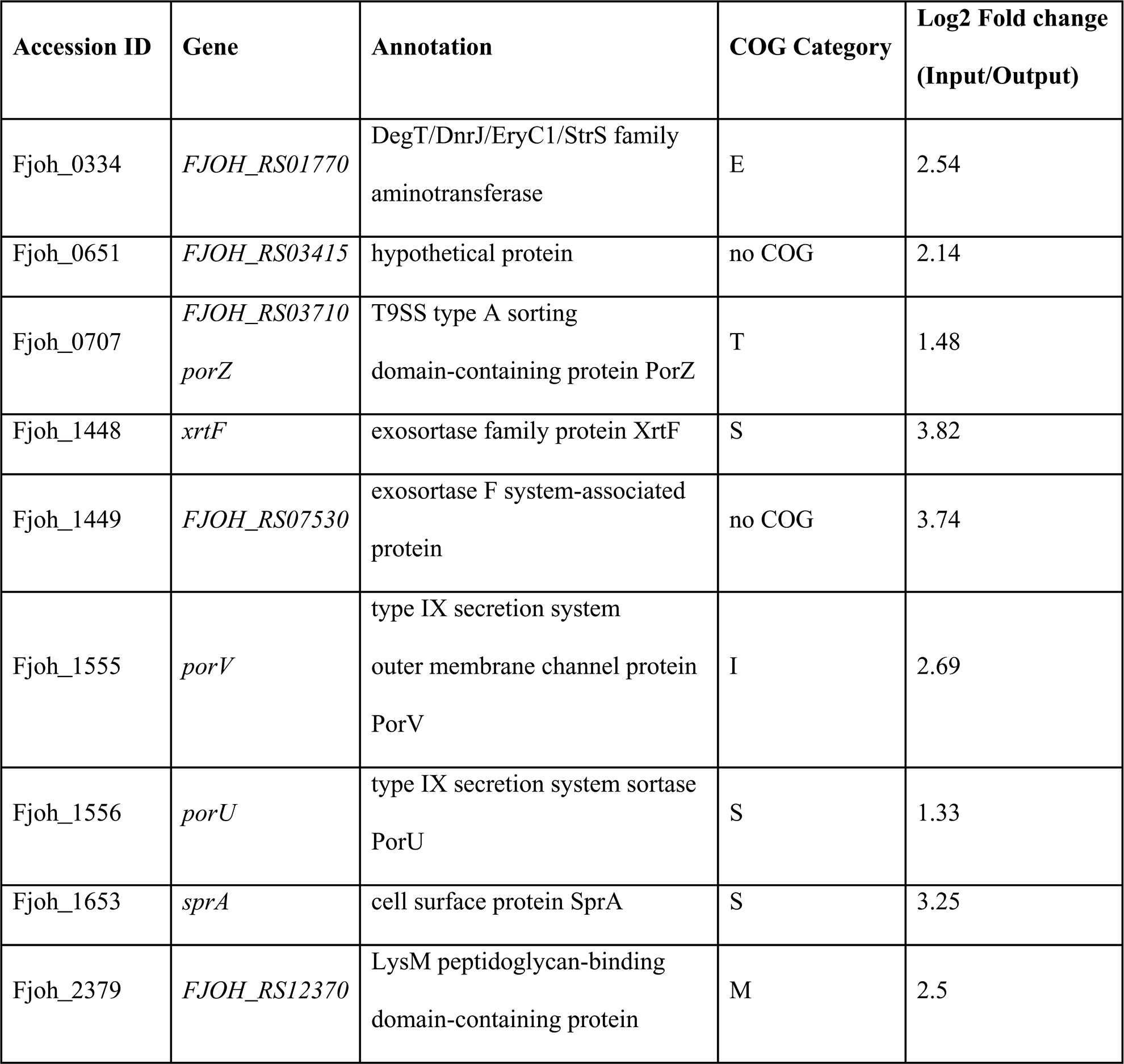
Sand colonization genes identified using INSeq screen and validated by mutant analysis. List of genes that were validated by constructing in-frame deletions. The complete list of genes identified is provided in Table S1. The COG categories are abbreviated as follows: E-amino acid metabolism and transport, T-signal transduction, S-unknown function, I-lipid transport and metabolism, M-cell wall/membrane/envelop biogenesis.

Functional annotation revealed that the largest group (40%) of candidate colonization genes had homologues in GenBank annotated as uncharacterized, encoding either hypothetical proteins or proteins of unknown function. A smaller group of genes (16%) encode proteins predicted to be involved in cell wall/membrane/envelope biogenesis, and other genes are predicted to be involved in transport and biosynthesis of primary and secondary metabolites (Fig. 1B).

We validated the INSeq screen by constructing in-frame deletions of nine genes that were underrepresented and assessed the mutants’ ability to colonize sand. Planktonic populations of the mutants in tryptic soy broth (TSB) were similar to those of the wild type after 48 h (Fig. 2A), whereas all nine mutants whose genes were underrepresented in the INSeq screen colonized sand poorly, indicating that the affected gene products contribute to sand colonization (Fig. 2B). Complementation of deletions with plasmid-borne copies of the gene of interest restored colonization to wild-type levels (Fig. S1).

**Figure 2:**
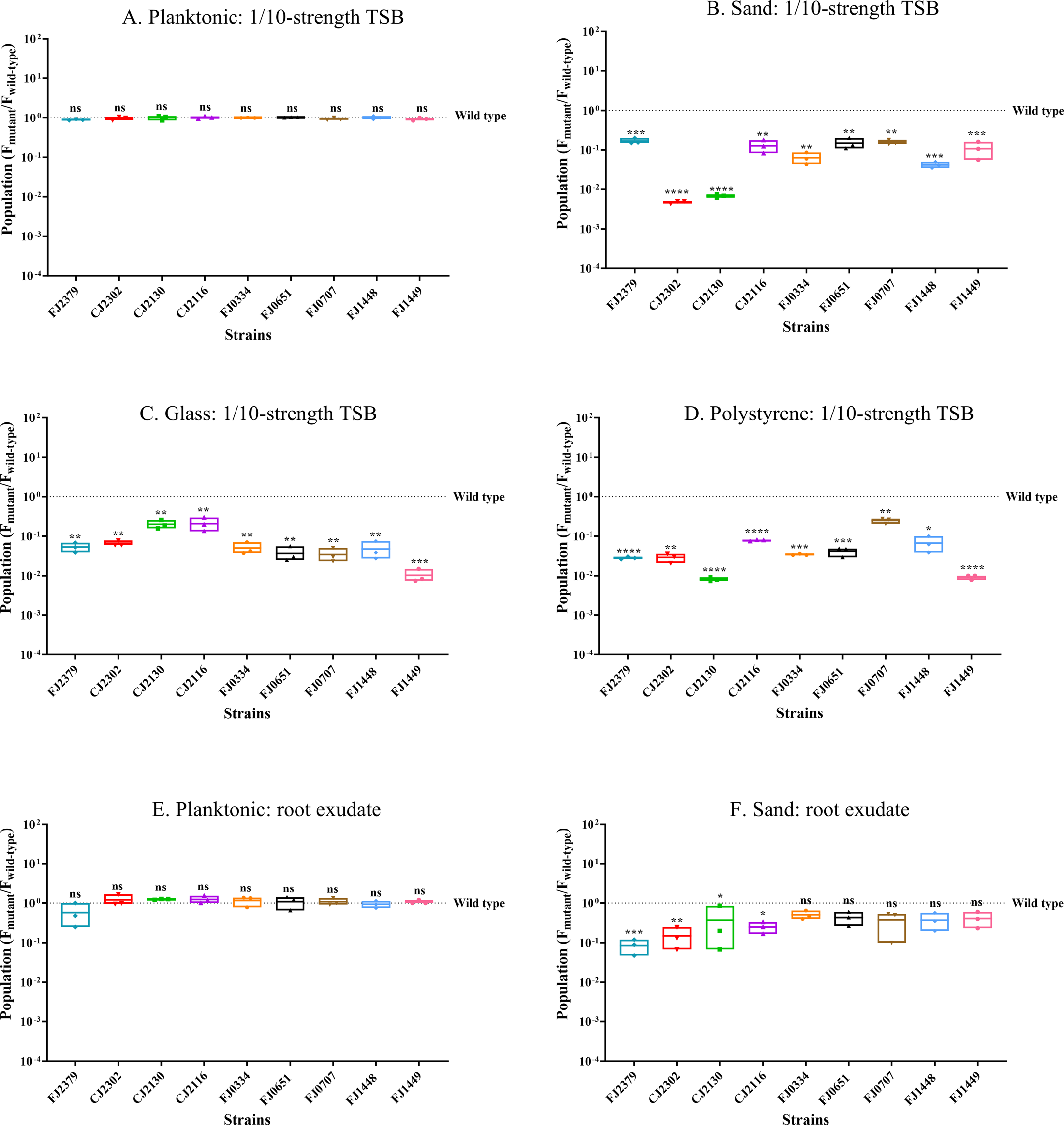
Surface colonization by mutants identified in INSeq analysis. Populations of wild-type *F. johnsoniae* CJ1827 and deletion mutants are represented as the ratio of mutant to the wild type. The initial inoculum was 10^6^ CFU/ml of wild-type *F. johnsoniae* CJ1827 or mutants (FJ2379, CJ2302, CJ2130, CJ2116, FJ0334, FJ0651, FJ0707, FJ1448, and FJ1449) and populations were determined after 48 h under the following conditions: (A) planktonic: 1/10-strength TSB, (B) sand: 1/10-strength TSB, (C) glass: 1/10-strength TSB, (D) polystyrene: 1/10-strength TSB, (E) planktonic: soybean root exudate, (F) sand: soybean root exudate. The dotted line indicates wild-type populations and each data point in the box plots represents one biological replicate. Statistical significance was evaluated using GraphPad Prism. Differences between the mutant and wild type are indicated as ns, not significant; *, p < 0.05; **, p < 0.01; ***, p < 0.001; ****, p < 0.0001.

Next, we determined whether the colonization phenotype was caused by a defect in attachment or the inability of the mutants to proliferate after attachment. We first quantified the population of *F. johnsoniae* CJ1827 (wild type) at several time points to investigate the influence of attachment and growth on colonization (Fig. S2). Based on our observations of the wild type, we measured the effects of gene deletions on attachment by assessing mutant populations during lag phase (T= 45 min) and the impact on growth by examining mutant populations during log phase (T=15 h) (Fig. 3). In the planktonic condition, most mutants (FJ2379, CJ2302, CJ2130, CJ2116, FJ0651, FJ0707, FJ1448, FJ1449) did not exhibit any significant difference in growth rate (Fig. S3) or population size. Mutant FJ0334 had a shorter doubling time than the wild type but reached stationary phase at a similar time to the wild type, thereby yielding a similar population size in the planktonic phase after 48 h. In contrast to the planktonic condition, eight mutants (CJ2302, CJ2130, CJ2116, FJ0334, FJ0651, FJ0707, FJ1448, FJ1449) exhibited poor colonization due to growth defect on sand, whereas only one mutant, FJ2379, displayed deficiencies in initial attachment. This suggests that the observed colonization phenotype is primarily driven by differences in the ability of these mutants to grow and establish themselves on the sand surface rather than their initial attachment capabilities.

**Figure 3:**
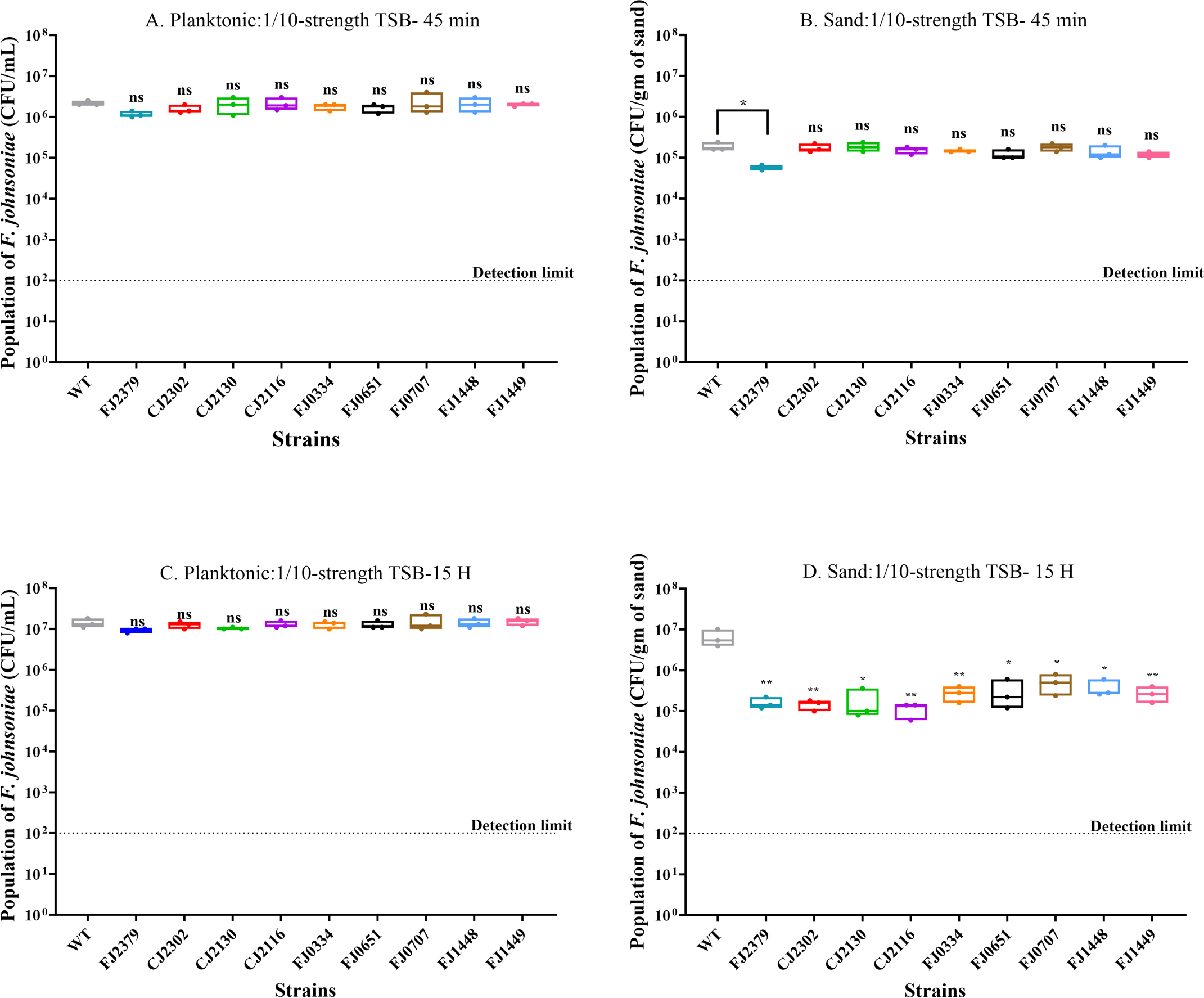
Impact of gene deletions on attachment and growth during sand colonization. Populations of wild type and mutants were determined after 45 min and 15 h under the following conditions: (A) 45 min no solid substrate (planktonic: 1/10-strength TSB) (B) 45 min sand: 1/10-strength TSB (C) 15 gh no solid substrate (planktonic: 1/10-strength TSB) (D) 15 H sand: 1/10-strength TSB. The dotted line indicates the detection limit (10^2^ CFU/mL) and each data point in box plots represents one biological replicate. Statistical significance was evaluated by comparing the mutants to the wild type with a one-way ANOVA followed by Dunnett’s test. Differences between the mutant and wild type are indicated as ns, not significant; *, p < 0.05; **, p < 0.01.

Additionally, since *F. johnsoniae* is a gliding bacterium, we determined whether any of these mutations affected motility (Fig. S4). We found that only one (CJ2302) of the nine mutants formed non-spreading colonies in PY2 agar, suggesting that not all mutations affecting colonization have an impact on motility.

### Sand colonization mutants are altered in colonization of other substrates

We tested the ability of the sand colonization mutants to colonize other substrates, such as glass, which is chemically similar to sand; polystyrene, which is structurally very different from sand; and sand in soybean root exudate to approximate the rhizosphere (Fig. 2). All nine mutants that colonized sand poorly were also defective in colonizing glass and polystyrene. This suggests that the deleted genes are important for colonization of surfaces generally. Deletion of genes mostly involved in cell wall/membrane biogenesis and T9SS led to poor sand colonization in root exudate, suggesting their importance in the rhizosphere. The other deletion mutants behaved similarly to the wild type on sand in root exudate.

### Biofilm formation correlates with surface colonization ability

Microorganisms colonize various surfaces in the soil, forming biofilms that enhance fitness by providing protection against predation, desiccation, exposure to antibiotics, and nutrient depletion (23–26). We tested the ability of colonization-defective mutants to produce biofilm using a crystal-violet assay (Fig. 4). All nine mutants that colonized poorly on sand, glass, and polystyrene produced less biofilm compared to the wild type, indicating that surface colonization is important for biofilm formation under these conditions. A linear regression analysis showed an association between biofilm formation and polystyrene colonization with a R^2^ value of 0.8632 (Fig. S5).

**Figure 4:**
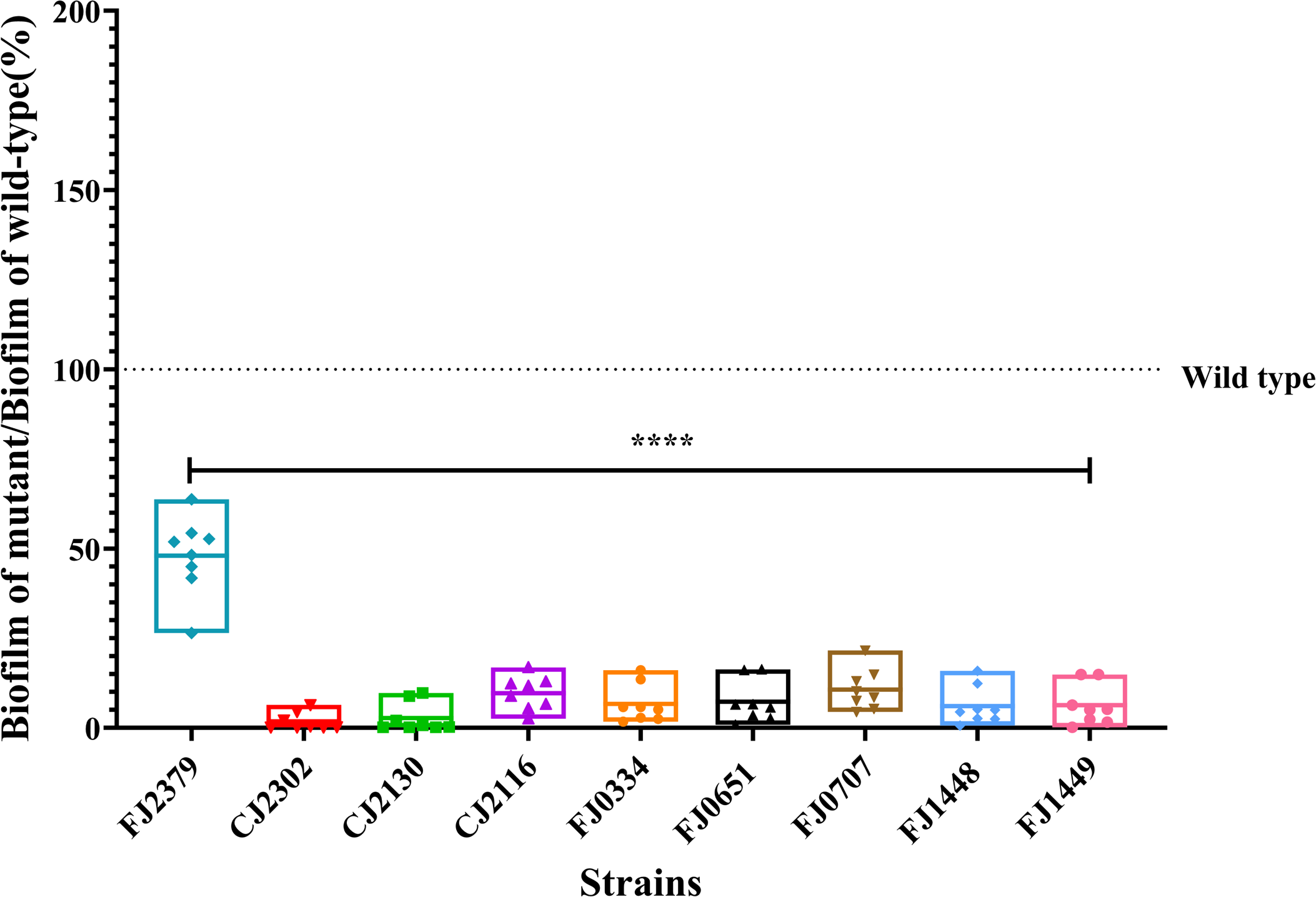
Colonization-defective mutants form less biofilm. Biofilm was quantified by crystal-violet staining (OD_595_) 18 h after inoculation. Each point represents one biological replicate comprising the data averaged from eight technical replicates. The data are expressed as a percentage of biofilm produced by the mutant relative to the wild type, where the dotted line indicates the level of wild-type biofilm. The OD_595_ values were square-root transformed to achieve equal variance, and statistical significance was evaluated by one-way ANOVA followed by Dunnett’s test. Differences between mutants and wild type are indicated as ****, p ≤0.0001.

### Surface colonization ability correlates with formation of zorbs

We previously reported that *F. johnsoniae* forms motile biofilms, designated zorbs, when observed in an under-oil microfluidic device (27). Zorbs move using cells at the base of the structure that are attached to the surface by one pole. Initially, the bacteria largely remain as single cells and then form microcolonies that migrate, merge, grow larger, and eventually disperse into single cells (27). To determine whether defects in static biofilm formation extended to motile biofilms, we monitored zorb formation in the mutants and wild type under oil on a glass substrate with bright-field time-lapse microscopy for 18 h (Fig. 5A) (Movie S1). In eight of the nine colonization-defective mutants (CJ2302, CJ2130, CJ2116, FJ0334, FJ0651, FJ0707, FJ1448, FJ1449), zorb formation was completely abolished. Interestingly, mutant FJ2379, which maps in a gene encoding a predicted LysM peptidoglycan-binding domain-containing protein, exhibited less zorbing capacity (the total area covered by zorbs over time) (Fig. 5B) and formed smaller zorbs than the wild type (Fig. 5C). These results indicate that genes required for normal surface colonization influence zorb formation and behavior.

**Figure 5:**
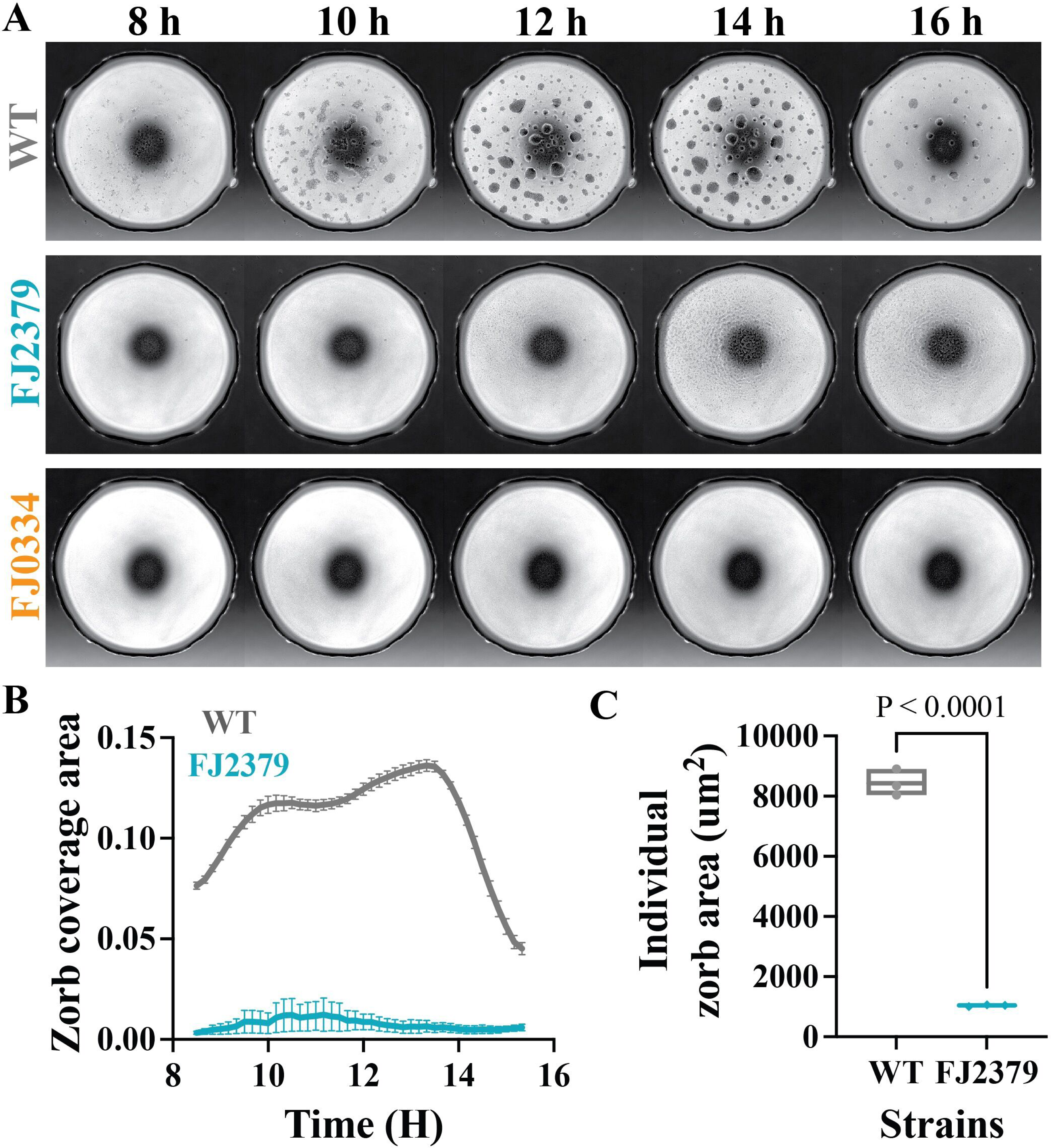
Colonization defects correlate with defects in zorb formation. Zorbing of wild-type and mutant strains was captured by brightfield imaging over 18 h and analyzed for differences in zorbing behavior. (A) Time-lapse images of wild type (CJ1827) and mutants (FJ2379, FJ0334 (representative of all eight mutants that did not form zorbs) between 8 and 16 h. (B) Plots of the total coverage area of zorbs given as a fraction of spot area over time. (C) Boxplot of time-averaged individual zorb area. Statistical significance was determined by a one-way ANOVA with multiple comparisons. All plots were generated by quantifying data from independent timelapse movies of three biological replicates for each strain.

### Colonization mutants exhibit altered behavior in a multispecies community

We used THOR to assess behavior of the surface colonization mutants in a community. THOR members exhibit several community-specific behaviors, including augmented expansion of *B. cereus* colonies induced by *F. johnsoniae* (10). Three mutants, CJ2130, CJ2302, and FJ2379, which contain deletions in genes encoding putative outer membrane proteins PorV, SprA, and LysM peptidoglycan-binding domain-containing protein respectively, induced more expansion of *B. cereus* colonies than the wild-type (Fig. 6). Initially, we hypothesized that expansion could be due to the relative difference in the level of biosurfactant produced by the mutants; however we did not observe any significant difference in the biosurfactant between the mutants and the wild type (data not shown). Given that PorV and SprA are integral to Type IX Secretion System (T9SS)-dependent secretion and anchoring proteins to the cell surface, deletion of *porV* and *sprA* may have resulted in the loss of secretion of specific cell-surface or outer-membrane proteins. This leads us to suggest that an interaction between surfaces of the two bacteria mediated by the outer membrane proteins in *F. johnsoniae* influences expansion of *B. cereus* colonies.

**Figure 6:**
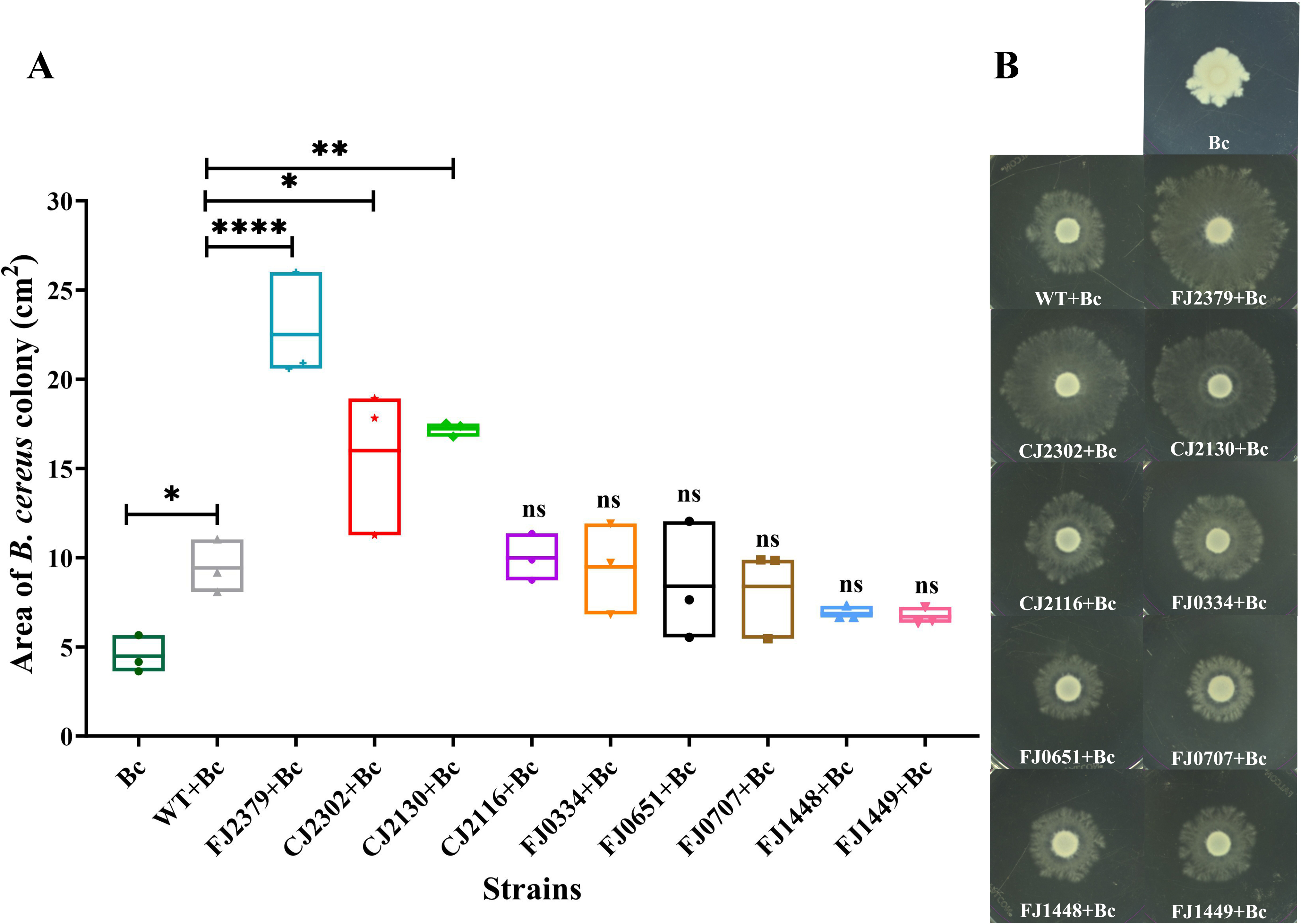
Colonization mutants affect *B. cereus* colony expansion. Colony expansion of *B. cereus* in 1/10^th^-strength TSA was observed in the presence of wild-type and mutant *F. johnsoniae* strains. The area of the *B. cereus* colony is shown as a box plot (A) and the experiment was performed three times and a representative image is shown in (B). The values (cm^2^) were square-root transformed for equal variance and the statistical significance was evaluated with a one-way ANOVA followed by Dunnett’s test. Differences between the mutants and wild type and between the wild type and *B. cereus* alone are indicated as ns, not significant; *, p ≤ 0.05; **, p ≤ 0.01, or ****, p ≤0.0001.

We next assessed whether surface colonization promotes fitness of *F. johnsoniae* in the community. In the absence of sand as a substrate (planktonic condition), the fitness of mutants defective in sand colonization was not altered in the full three-member community, as indicated by the mutants and the wild-type achieving similar populations in the community. The one exception was FJ2379, whose populations were lower (Fig. 7A) than the wild type in the planktonic community. Consequently, we examined the impact of cell-free supernatants from *P. koreensis*, *B. cereus*, and both *P. koreensis* and *B. cereus* on the wildtype and FJ2379. We found that whereas FJ2379 did not exhibit any growth defect when cultured in its own or in the culture supernatant of *B. cereus*, it achieved a significantly lower population when cultured in the culture supernatants of *P. koreensis* and *P. koreensis*+*B. cereus* (data not shown). This suggests that the observed mutant phenotype in the planktonic community is likely mediated by metabolites secreted by *P. koreensis,* and FJ2379 appears to be relatively sensitive to these metabolites. In contrast, all mutants defective in sand colonization in solitary culture survived poorly in the full community compared to the wild type on sand (Fig. 7B), indicating that the genes required for solitary colonization of surfaces are also required for *F. johnsoniae* success in a microbial community on a surface.

**Figure 7:**
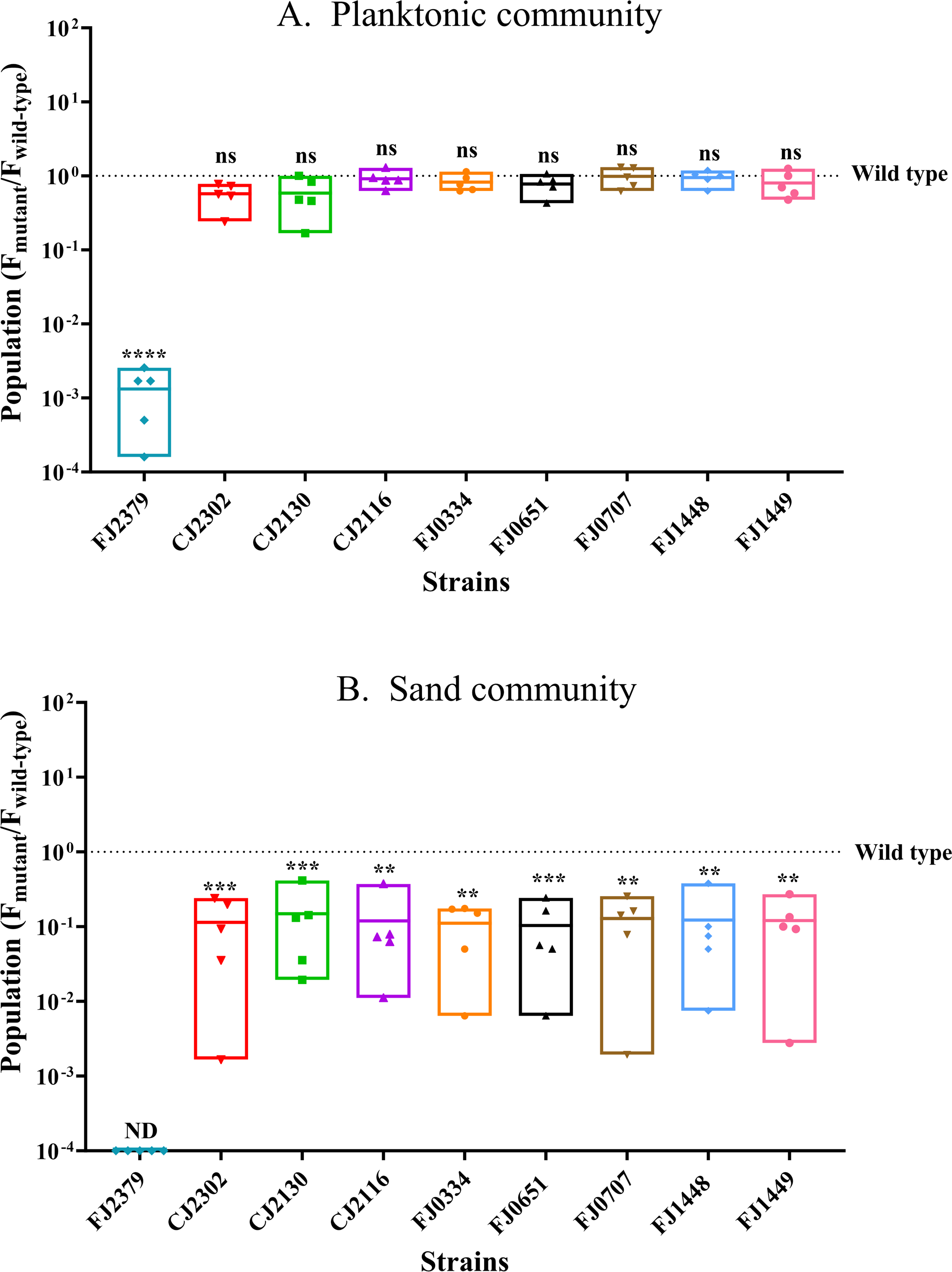
Colonization ability affects fitness of *F. johnsoniae* in THOR. (A) *F. johnsoniae* wild type and mutants when grown with *P. koreensis* and *B. cereus* planktonically. (B) *F. johnsoniae* wild type and mutants when grown with *P. koreensis* and *B. cereus* in the presence of sand. Each data point represents one of five biological replicates, with the dotted line indicating the value for the wild type. The CFU values were log_10_ transformed for equal variance. The statistical significance was evaluated by comparing the mutants to the wild type with a one-way ANOVA followed by Dunnett’s test. ns = not significant; **, p ≤ 0.01; ***, p ≤ 0.001; or ****, p ≤ 0.0001.

## DISCUSSION

In this study, we used INSeq, a genetic screen that couples transposon mutagenesis and high-throughput sequencing, to identify genes in *F. johnsoniae* that are important for sand colonization in solitary culture and in a model community. INSeq [highly similar to transposon sequencing (TnSeq)] is a powerful genetic tool to establish relationships between genes and bacterial behavior (28, 29). This tool has been used to simultaneously evaluate the fitness of thousands of discrete mutants and has identified genes crucial for root colonization in several Proteobacteria and genes related to soil colonization in others (30–34). Although INSeq technology was developed in *Bacteroides thetaiotaomicron,* the work presented here is the first INSeq screen on another member of the Bacteroidetes phylum to reveal genes necessary for sand colonization.

Sand is an abundant soil particle, and the presence of *F. johnsoniae* promotes aggregation of sand particles, thereby reducing soil erosion (35). We identified 25 genes related to sand colonization, most of which are either uncharacterized or encode proteins with unknown functions. Other gene categories found in the screen were annotated as involved in cell wall/membrane/envelope biogenesis and transport and metabolism of primary and secondary metabolites (Fig. 1). Although the majority of these genes have not been explored thoroughly in *F. johnsoniae*, their predicted functions are similar to genes previously shown to play roles in fitness of other rhizosphere-colonizing bacteria (30, 31).

We validated the INSeq screen by constructing in-frame deletions in nine genes and showed that the mutants’ abilities to colonize various substrates, form biofilms, and survive in a model rhizosphere community were affected by the deletions (Table 2). Based on the predicted functions of the deleted genes, these mutants can be divided into four categories: Type IX secretion system (T9SS), exosortase-system, cell-surface components, and hypothetical proteins.

**Table 2:**
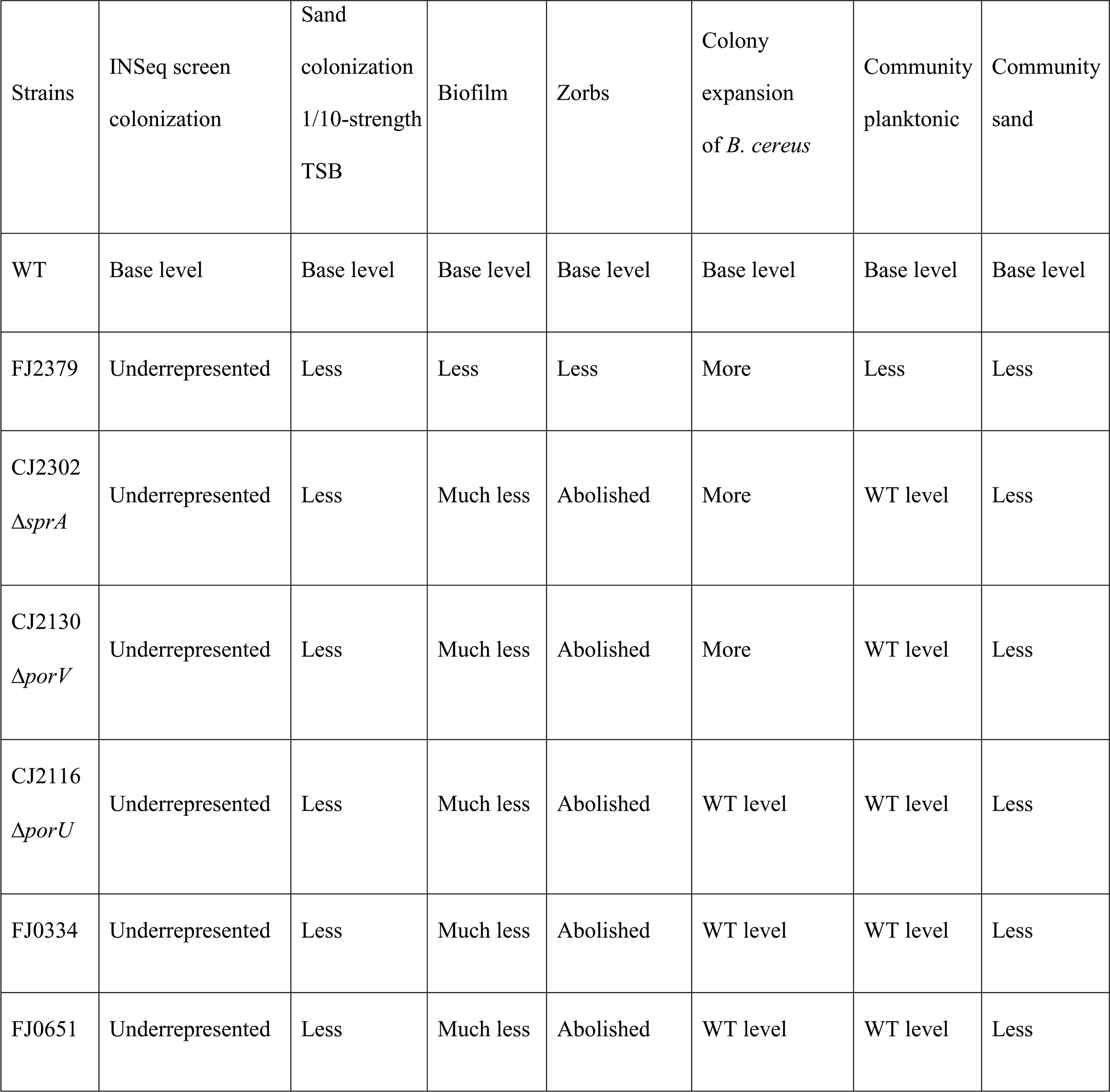
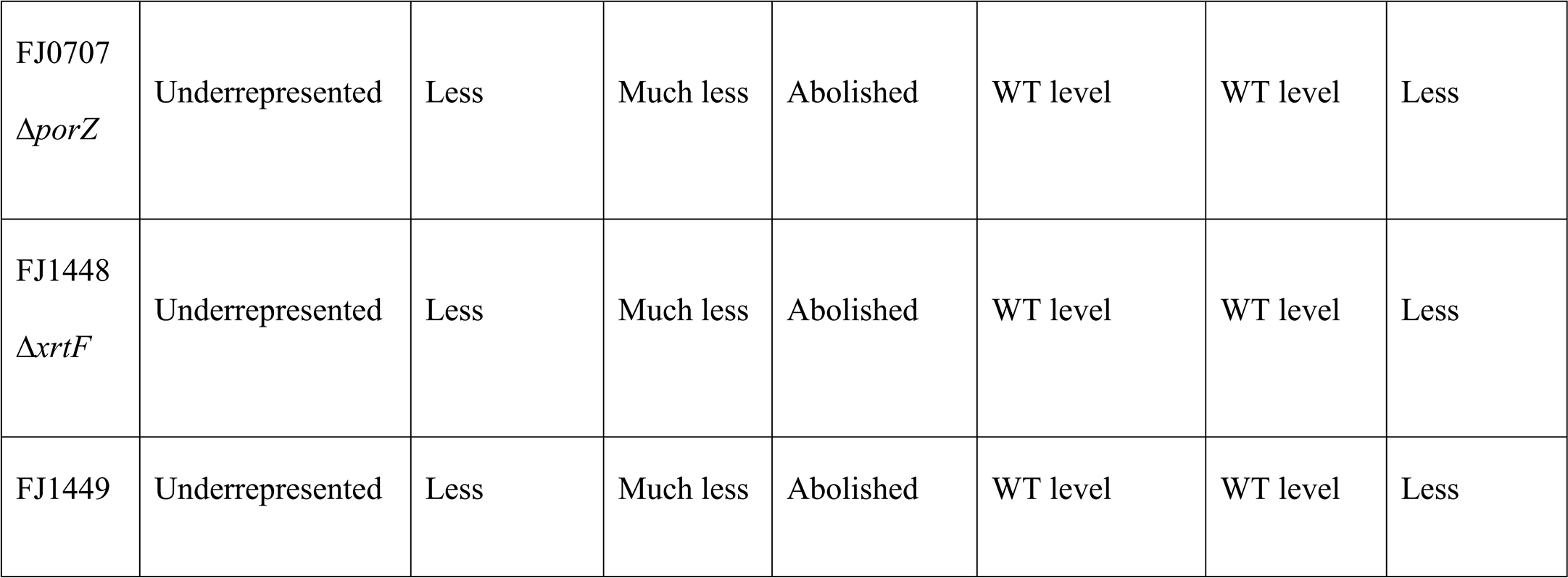
Summary of surface-colonization mutant phenotypes.

Numerous studies of non-gliding bacteria have demonstrated the importance of motility appendages such as flagella and pili in colonization of surfaces in the rhizosphere (36–39). In contrast, *F. johnsoniae* glides over surfaces without the aid of flagella or pili, instead using gliding motility apparatus and adhesins secreted by T9SS (15). Deletion of genes involved in T9SS or gliding motility have been shown previously to affect attachment to glass (*sprA*, *sprE, gldK*, *gldL*, *gldM, porV*) and plant roots (*gldJ*) and biofilm formation (*gldA*, *gldB*, *gldD*, *gldF*, *gldG*, *gldH*, *gldI*, *gldJ*, *gldN* and *porV*) (40–45). Our study adds to the existing body of research by demonstrating that deletion of genes associated with T9SS (*FJOH RS03710* [*porZ* orthologue of *Porphyromonas gingivalis* W83], *porV*, *porU*, and *sprA*) does indeed reduce colonization of various substrates and abolishes biofilm formation. Proteins PorV, PorU, and PorZ are integral components of the T9SS cell surface-exposed attachment complex (PorQUVZ), with PorV playing a pivotal role in shuttling cargo proteins from the SprA translocon to the attachment complex (46–48). Subsequently, these cargo proteins are processed and anchored to the cell surface. Given that the proteins encoded by these genes play a crucial role in the T9SS-mediated secretion of cell-surface and extracellular proteins, deletion of these genes most likely caused loss of secretion of many cell surface proteins or adhesins resulting in the observed defects in surface colonization and biofilm formation.

Several genes from the INSeq screen involved in sand colonization also affected the formation of motile biofilms, or zorbs, produced by *F. johnsoniae*. Zorbs are self-propelled spherical microcolonies with an EPS core that move using cells at the base of the zorbs (“base cells”) that attach to the surface by one pole of the cell (27). Our data genetically link surface attachment and zorb formation, confirming that the initial attachment of base cells to the surface is necessary for zorb formation. This study also demonstrates the relatedness of stationary and motile biofilms by showing that zorbing was absent in mutants that failed to produce any detectable stationary biofilm. In contrast, mutant FJ0347, which produced more biofilm, formed hyper-motile zorbs that merged more than the wild type (Fig. S6).These findings corroborate previous work showing that zorbs are authentic biofilm structures.

We identified two genes (*xrtF* and *FJOH_RS07530*) predicted to encode exosortase family proteins. Exosortases are proteases that cleave proteins that contain the carboxy-terminal protein-sorting signal PEP-CTERM and mostly occur in bacterial species with exopolysaccharide biosynthesis gene clusters (49, 50). Although exosortases have been proposed to be associated with biofilm formation based on bioinformatic analysis, our study is the first to provide empirical data indicating that exosortases play a role in surface colonization and biofilm formation. Future research will focus on the substrates for exosortases in *F. johnsoniae*, which are predicted to be quite dissimilar from PEP-CTERM signal sequences based on genomic analysis.

We additionally identified four colonization genes within a 76.5-kb gene cluster that is predicted to synthesize and export cell-surface polysaccharides (51). This gene cluster contains several transcriptional units, and deletion of genes from various regions of it had diverse effects on biofilm formation. For instance, deletion of gene *FJOH_RS01770,* which is part of a transcriptional unit predicted to be involved in biosynthesis of dTDP-β-L-rhamnose and fucose, abolished biofilm formation, whereas deletion of gene *FJOH_RS01835,* predicted to be involved in LPS biosynthesis, enhanced biofilm formation (Fig. S6A). This difference in phenotype between the two mutants is likely due to FJ0347 having a more hydrophobic cell surface, a characteristic that contributes to adhesion and biofilm formation (Fig. S6B) (27, 52–54).

We predicted that genes that are uniquely involved in interactions between *F. johnsoniae* and the soil environment would be enriched for those of unknown function since the majority of functional characterization of bacterial genes has been conducted in pure culture in laboratory media. The data support this prediction and demonstrate the power of ecological screens to increase the functional assignments of previously unknown genes. Of five genes (*FJOH RS02845*, *FJOH RS03290*, *FJOH RS03415*, *FJOH RS0536*, and *FJOH RS15790*) identified in the INSeq screen that encode hypothetical proteins, we established the role of *FJOH RS03415* in sand colonization and biofilm formation.

The study lays the groundwork for understanding *F. johnsoniae* soil colonization and success in a community. It also suggests several themes to test more broadly among rhizosphere bacteria. First, we provide evidence that the same genes that are required for surface colonization by *F. johnsoniae* in solitary culture are also important for its survival in community, a connection not previously addressed with genetic analysis in rhizosphere bacteria. Second, we provide further evidence that cell-surface molecules, such as proteins and polysaccharides, are significant actors in *F. johnsoniae* surface colonization, which has been established previously in other bacteria. This finding is especially important because of the role of these same molecules in community fitness. Finally, identifying six genes of unknown function in the colonization screen suggests that ecological screens can identify functions for genes that do not play substantial roles in standard laboratory pure culture conditions, which comprise the majority of previous genetic screens in microbiology. Similar work with other species will establish whether these themes are universal principles in microbial ecology.

## MATERIALS AND METHODS

### Bacterial strains, plasmids, and growth conditions

Strains and plasmids used in this study are listed in Table 3 and primers are listed in Table S2. Antibiotics were used at the following concentrations when needed: ampicillin 100 μg ml^-1^; erythromycin 100 μg ml^-1^; kanamycin 50 μg ml^-1^; streptomycin 100 μg ml^-1^; and tetracycline 20 μg ml^-1^. Bacteria were propagated on 1/2-strength tryptic soy broth (TSB) at 28°C with shaking. For experimental setup, 10^6^ or 10^7^ cells of each bacterium were inoculated from saturated overnight cultures and grown statically in 1/10-strength TSB at 20°C. Populations of these strains were quantified by dilution plating on LB agar supplemented with gentamicin (10 μg ml^-1^) for *F. johnsoniae*, ampicillin (100 μg ml^-1^) and erythromycin (5 μg ml^-1^) for *P. koreensis*, and polymyxin B (5 μg ml^-1^) for *B. cereus*. *Escherichia coli* strains used for cloning were grown in Luria–Bertani broth (LB) at 37°C.

**Table 3:**
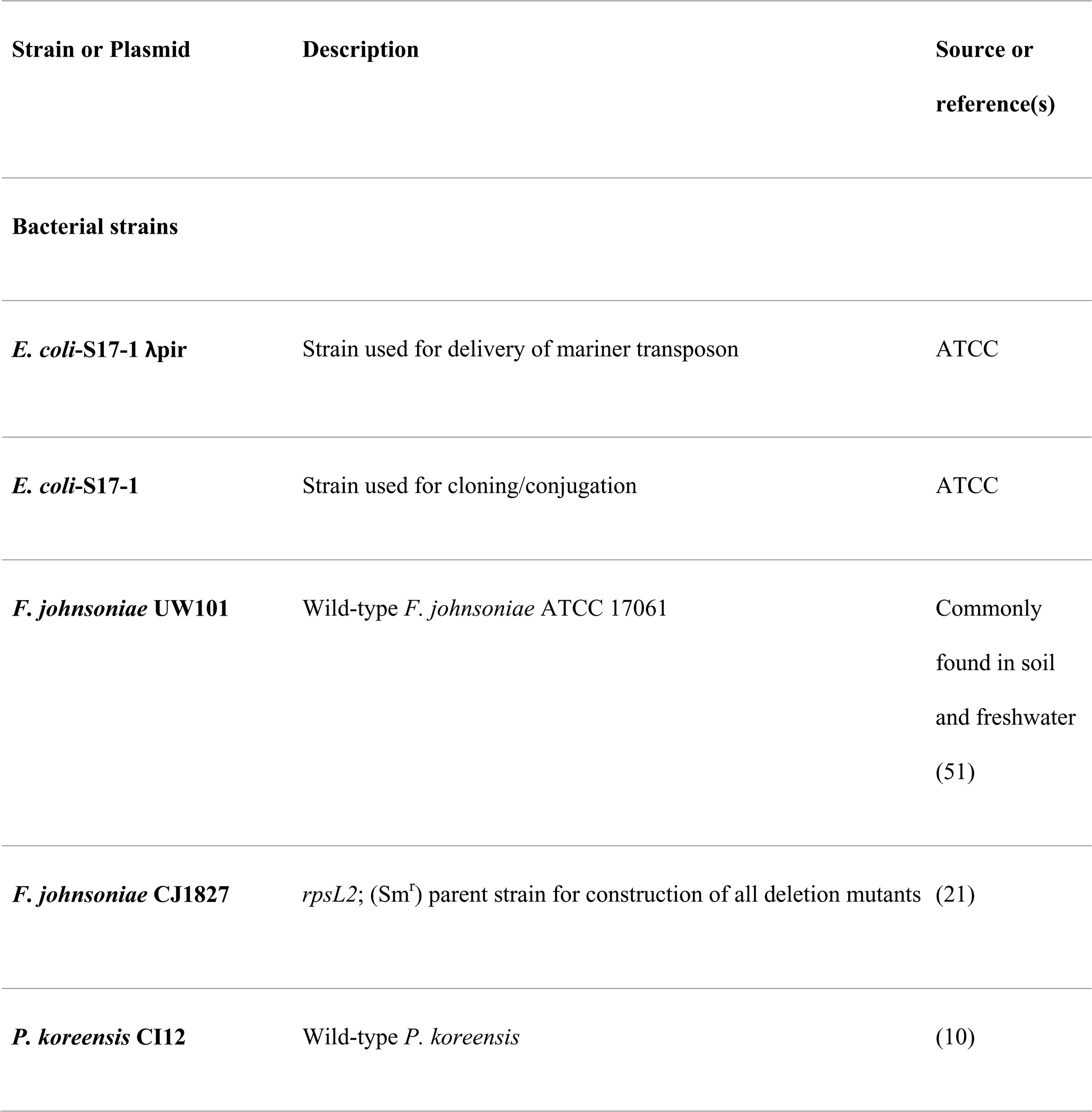
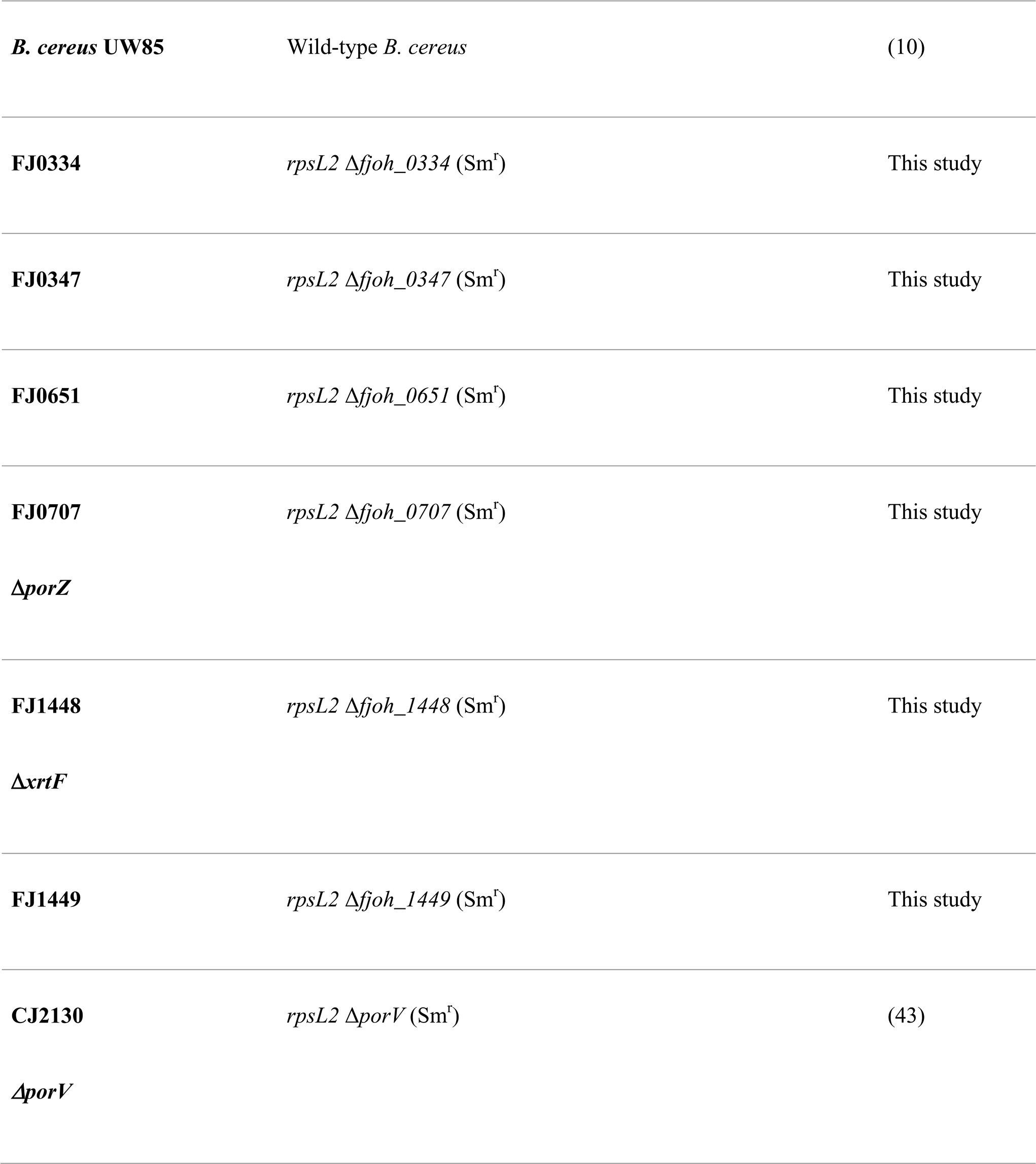
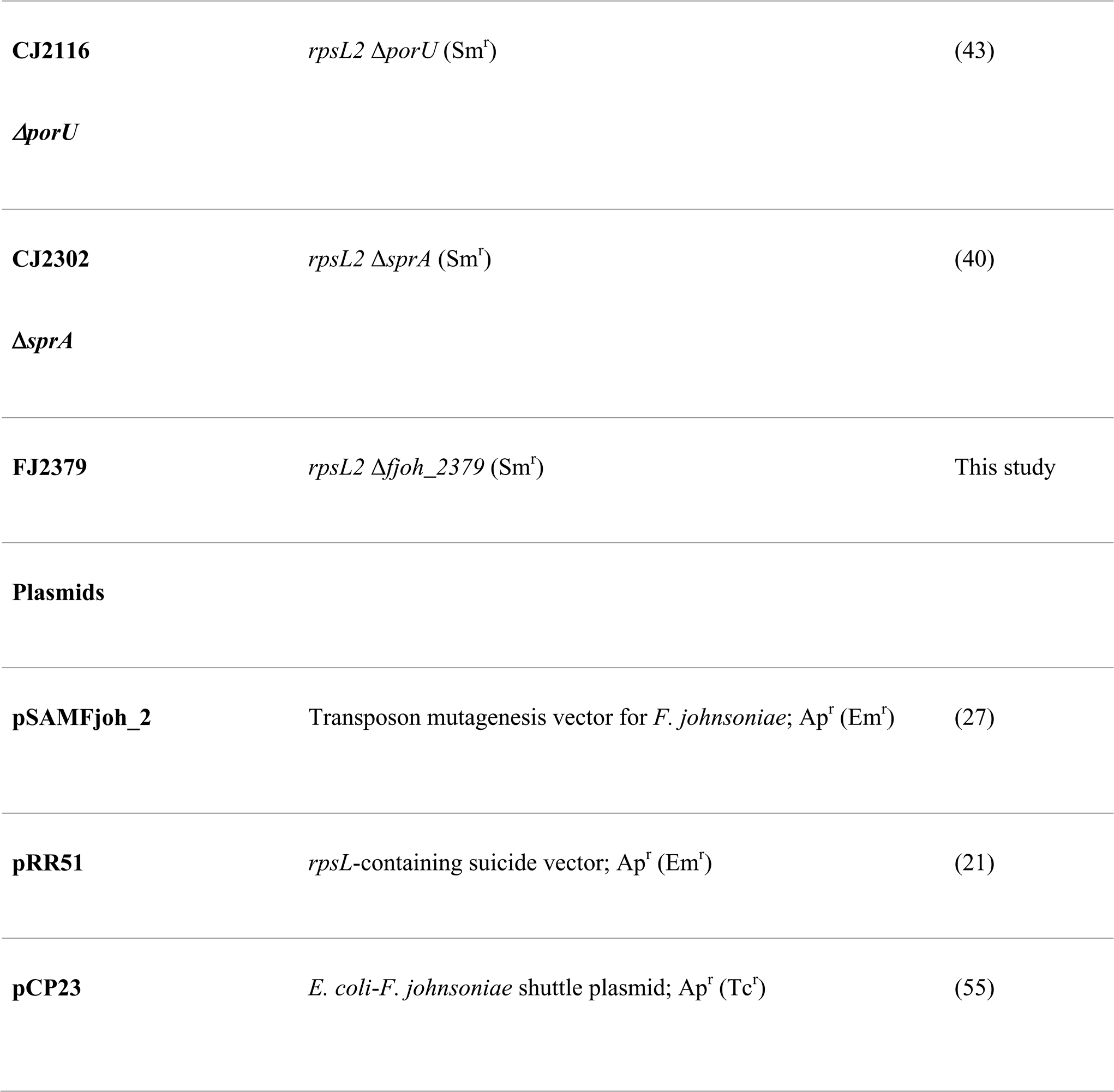
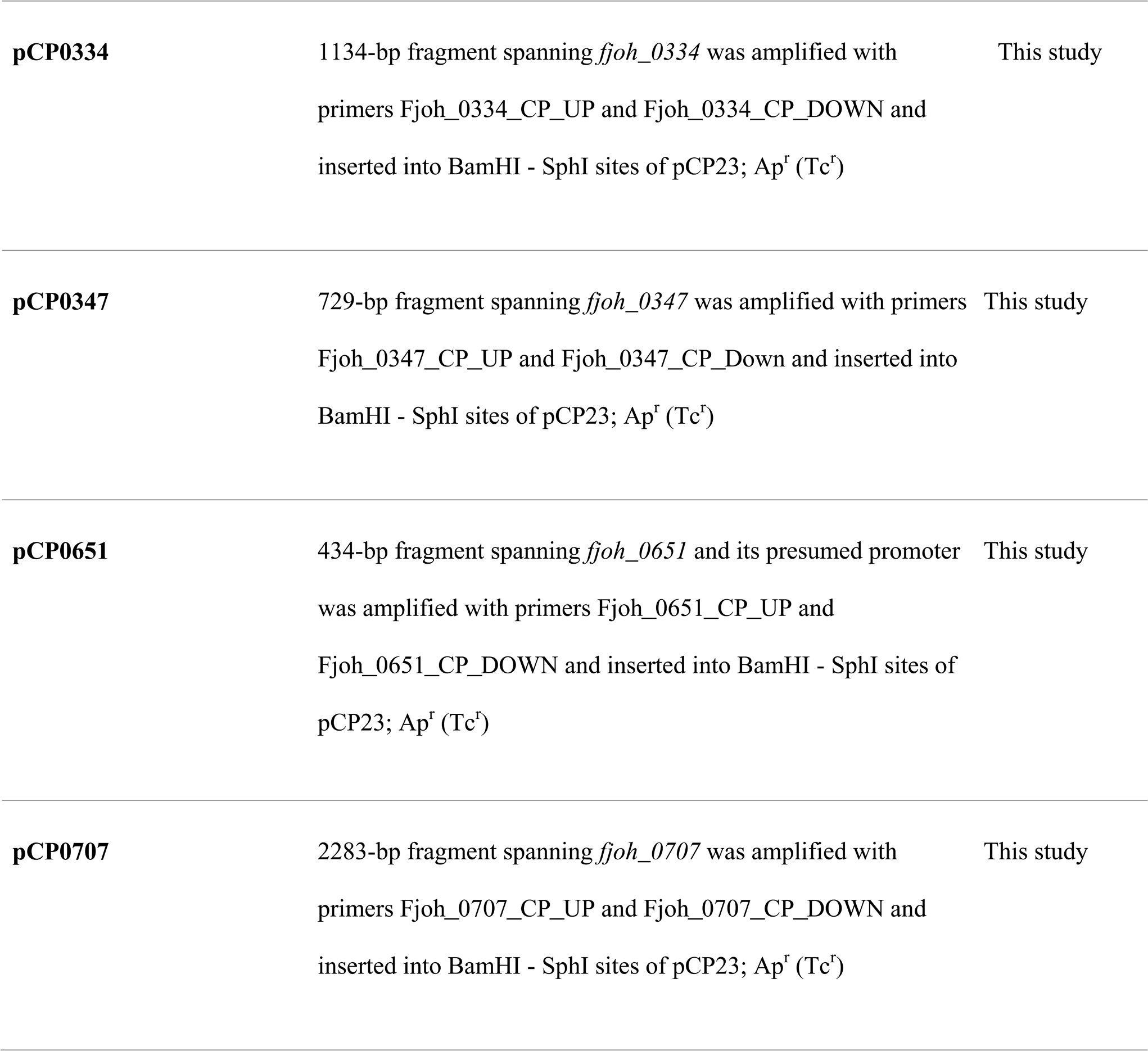
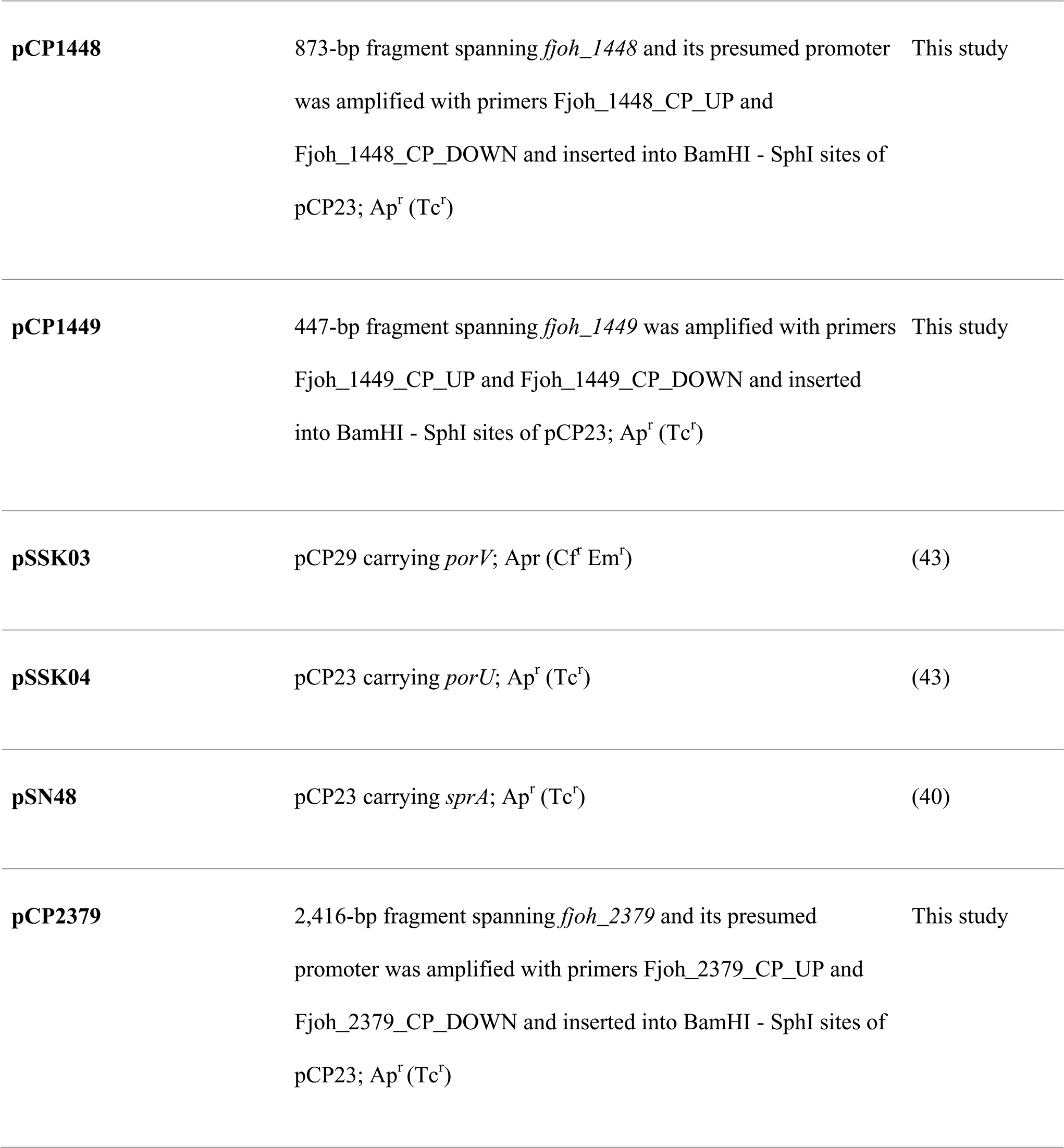
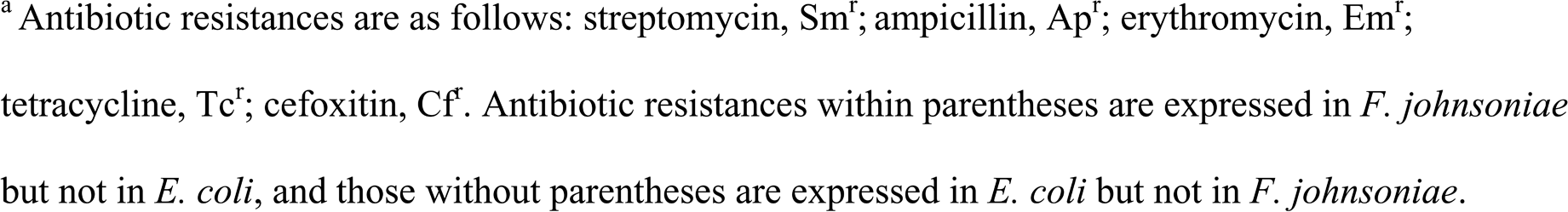
Bacterial strains and plasmids used in this study.

### Construction of mariner transposon for *F. johnsoniae*

The *F. johnsoniae* mariner transposon delivery vector pSAMFjoh_2 was constructed by modifying the resistance cassette and transposase promoter in the *B. thetaiotaomicron* pSAM-BT42 vector as previously described (27). Briefly, it was modified by replacing the *B. thetaiotaomicron* transposase promoter with the *F. johnsoniae rpoD* promoter and adding an *F. johnsoniae-*compatible erythromycin-resistance (erm) cassette.

### Mutant library construction

The transposon mutant library was constructed by conjugating *F. johnsoniae* UW101 with *E. coli*-S17-1 λpir containing the MmeI-adapted mariner transposon in pSAM_Fjoh2. Briefly, the donor (*E. coli* S17-1 λpir) and recipient (*F. johnsoniae* UW101) strains were grown overnight in LB supplemented with ampicillin (100 μg ml^-1^) and in casitone yeast extract (CYE) media respectively. Cultures were aliquoted in a 1:4 donor: recipient ratio, centrifuged, and washed with 1 ml CYE medium. The cells were then resuspended in 100 µl of CYE and spotted on CYE agar. Conjugation took place at 28°C for 12 h. Exconjugants were selected by plating on CYE medium supplemented with erythromycin (100 μg ml^-1^) and kanamycin (50 μg ml^-1^) for counterselection against the donor *E. coli* strain. The plates were incubated at 28°C for 48 h, then ∼75,000 clones were pooled into fresh CYE, and aliquots of the mutant library containing 40% glycerol were stored at -80°C until further use.

### System for INSeq selection in *F. johnsoniae*

Overnight cultures of the *F. johnsoniae* mariner transposon library were made from aliquots of the mutant library in 1/2-strength TSB. Cells were pelleted, washed, and resuspended in 5 ml of 10 mM NaCl. One million cells (OD_600_ = 0.0008) from the *F. johnsoniae* mariner transposon library were inoculated in 1 ml of 1/10-strength TSB with and without 0.5 g of sterile sand (50-70 mesh particle size; Sigma-Aldrich), and then incubated without shaking at 20°C for 48 h. The tubes with sand were washed three times with 10 mM NaCl to remove loosely adhered cells and resuspended in 1 ml of 10 mM NaCl. All the tubes were vortexed and sonicated prior to DNA extraction. The QIAGEN DNeasy blood and tissue kit was used to extract DNA, which was used as a template to amplify mariner transposon insertion sites and was ligated with appropriate barcodes as previously described (28). The 125-bp amplicons were adjusted to 1 nM and sequenced using the Illumina HiSeq platform with 2×150-bp PE configuration.

### INSeq data analysis

Pooled raw sequencing reads were quality processed with fastp [https://doi.org/10.1093/bioinformatics/bty560) using --disable_adapter_trimming and --cut_tail (QC trimming at the 3’ end only) parameters] and then split into individual fastq files (one for each replicate) by their indexed adapter sequence. Reads were aligned to *F. johnsoniae* UW101 (accession GCF_000016645.1 ASM1664v1) with bowtie2 (https://www.nature.com/articles/nmeth.1923) using default parameters, and mapped reads were matched to corresponding genes based on *F. johnsoniae* UW101 coding-sequence annotations (same accession as above). Counts per million (CPM) were calculated in edgeR (https://academic.oup.com/bioinformatics/article/26/1/139/182458).

### Construction of deletion mutants and complementation plasmids

In-frame deletions in genes *FJOH_RS01770, FJOH_RS01835, FJOH_RS03415, porZ, xrtF, FJOH_RS07530, porV, porU, sprA, FJOH_RS12370* were constructed using a gene-deletion strategy that uses dominance of wild-type *rpsL* on the plasmid pRR51 as a counter-selectable marker over the chromosomal mutant *rpsL* in *F. johnsoniae* CJ1827 (21). Briefly, primers were designed with an overlapping sequence of 18 bp to amplify and fuse the regions upstream and downstream of the gene of interest using SOE-PCR. The 2-kb fusion product was ligated to vector pRR51 and was transformed into *E. coli* S17 by electroporation. *E. coli* S17 cells carrying pRR51 with the 2-kb insert were conjugated with *F. johnsoniae* CJ1827, and exconjugants were selected using CYE supplemented with 100 μg ml^-1^ erythromycin. Erythromycin-resistant (streptomycin-sensitive) clones were grown overnight in CYE without antibiotics and plated on CYE agar medium supplemented with streptomycin 100 μg ml^-1^ to select for the loss of plasmid by a second recombination event. Deletion of the target gene in *F. johnsoniae* CJ1827 was confirmed by PCR amplification using upstream forward and downstream reverse primers and sequencing of the resulting 2-kb product. The wild-type strain used in all mutant comparisons is *F. johnsoniae* CJ1827 unless otherwise specified.

The plasmids used for complementation were derived from pCP23. The gene of interest with its presumed promoter was amplified using their respective primers. Both the insert and the vector were digested with BamHI-HF and SphI-HF and were ligated into their respective sites in pCP23. The genes that were in a cluster or did not have their own promoter were amplified and cloned into BamHI-HF and SphI-HF sites of pCP23 in the direction of orf1 and were expressed by the vector promoter.

### Surface colonization experiments

Surface colonization was assessed using three substrates: sand (0.5 gm; 50-70 mesh particle size; Sigma-Aldrich), glass (2 gm, 3 mm bead diameter; Fisher Scientific) and polystyrene (24-well plates; CELLTREAT Scientific Products). The wild type (*F. johnsoniae* CJ1827) and mutants were grown individually for 20 h in 1/2-strength TSB at 28°C; 5 ml of the overnight culture was centrifuged, washed once, and resuspended in 10 mM NaCl. The wild type and mutants (10^6^ CFU ml^-1^) were inoculated individually in 1/10-strength TSB with and without substrates and grown for 48 h statically at 20°C. The samples were washed three times with 10 mM NaCl to remove loosely adhered cells and were vortexed 30 s, sonicated (Bransonic ultrasonic cleaner 1510R-MT) gently for 2 min with ∼50Hz, and vortexed 30 s to retrieve cells adhered to the substrates. This method was effective in detaching most colonized cells without clumping or lysing them. For polystyrene, each well was scraped 100 times using a sterile toothpick instead of vortexing before and after sonication. The population of cells adhered to the surface was determined by serial dilution and plating on LB containing gentamicin (10 μg ml^-1^). The soybean root exudate for surface colonization experiments were made as previously described (10). Colonization experiments with planktonic, sand, glass, and polystyrene conditions with 1/10-strength TSB were each performed individually three times with one mutant and wild type at a given time point. We compared each mutant to its wild type by analyzing log_10_ transformed CFU values using an unpaired two-tailed t- test with Welch’s correction to account for equal variance. Experiments with root exudate were performed three times with all ten mutants and the wild type. Hence, we log_10_ transformed the values to achieve equal distribution and performed a one-way ANOVA followed by Dunnett’s test.

### Biofilm quantification

*F. johnsoniae* biofilm formation was quantified using crystal-violet staining. First, 150 μl of sample containing 10^7^ cells ml^-1^ of *F. johnsoniae* strains in 1/10-strength TSB was inoculated into each well in a 96-well plate and incubated without shaking at 20°C for 18 h. The planktonic cells were discarded, and the plate was washed with water. The biofilm was stained with 200 μl of 0.1% crystal-violet solution in 20% methanol for 15 min and washed three times with water. Then 33% acetic acid (200 μl/well) was used to solubilize the crystal violet and the absorbance was measured at 595 nm.

### Zorb microscopy and analysis

An under-oil open microfluidic system (UOMS) device was constructed by chemical vapor-deposition of polydimethylsiloxane (PDMS)-silane onto a glass substrate. The PDMS-silane grafted surface was patterned by O_2_ plasma treatment to generate patterns of treated and non-treated areas as previously described (56). The device was overlaid with 1.5 mL Fluorinert FC-40 oil and UOMS spots were seeded with 10^7^ cells (2 µL) of wild type or mutant *F. johnsoniae* in 1/10-strength TSB. The UOMS device was observed with bright field time-lapse microscopy for 18 h at room temperature as previously described (27).

Time-lapse movies were analyzed using a custom MATLAB code to threshold each frame using Otsu’s segmentation method, then extracting the boundary of each zorb to obtain measures of the zorb area. Zorb speed was quantified using MATLAB code from a previously described tracking method (57) from videos taken at 30 min intervals. Zorb merging was measured manually by counting the number of merging events of individual zorbs over a period of 8 h (to include zorb formation and subsequent dispersion). Four zorbs were randomly selected and tracked throughout a given time lapse (one in each of the four quadrants to ensure independent zorbs were tracked), with a total of three time-lapse movies analyzed for each strain, resulting in 12 replicates for each strain.

### *B. cereus* motility assay

The expansion of *B. cereus* colonies in the presence of wild-type or mutant *F. johnsoniae* was tested using the previously described modified spread-patch method (10). Briefly, *B. cereus* and *F. johnsoniae* strains were grown individually for 20 h at 28°C, and 1 ml of culture of each strain was pelleted at 6,000 x *g* for 6 min and then resuspended in 1 ml of PBS. Wild-type or mutant *F. johnsoniae* culture (100 μl) was spread on 1/10-strength TSA plates, which were dried for 2 h at 28°C. *B. cereus* culture (10 μl) was spotted at the center of each plate and incubated at 28°C. The plates were monitored and imaged on day 5.

### Determining fitness of *F. johnsoniae* in THOR

Fitness of *F. johnsoniae* in THOR was determined by growing 10^7^ cells ml^-1^ of *F. johnsoniae* strains with equal CFU of *B. cereus* and *P. koreensis* (1:1:1) statically for 48 h in 1/10-strength TSB with and without sand at 20°C. The samples were processed as described for the surface colonization experiment and the populations of *F. johnsoniae, P. koreensis,* and *B. cereus* were quantified by plating the sample on LB agar supplemented with gentamicin (10 μg ml^-1^), ampicillin (100 μg ml^-1^) plus erythromycin (5 μg ml^-1^), and polymyxin B (5 μg ml^-1^) respectively.

## DATA AVAILABILITY

The data are available from the corresponding author. Bacterial strains and mutant libraries are available upon request. The primers used in this study are listed in Table S2. Representative zorb videos can be found in the supplemental material and the rest will be made available upon request. Insertion sequencing data can be found in the National Center for Biotechnology Information’s Sequence Read Archive under BioProject accession PRJNA955258. The files Ama01, Ama02, and Ama03 correspond to three biological replicates in the planktonic condition and the files Ama04, Ama05, and Ama06 correspond to three biological replicates in the sand condition.

## ACKNOWLEDGMENTS

This study was supported by the U.S. Army Research Laboratory and the U.S. Army Research Office under contract/grant W911NF1910269, the Vilas Trust, the Office of the Vice Chancellor for Research and Graduate Education at the University of Wisconsin-Madison, the University of Wisconsin Carbone Cancer Center Support Grant NIH P30CA014520, and NIH R01AI59940. A. Hurley was supported by USDA NIFA Grant 2019-2018-08058 (Accession no. 1019190). J.F. Nepper was supported by an NHGRI training grant from the Genomic Sciences Training Program 5T32HG002760. M.G. Chevrette was supported by USDA NIFA Grant 2020-67012-31772 (Accession no. 1022881). C. Li was supported by NSF EFRI-1136903-EFRI-MKS, NIH R01 CA247479, NIH R01 AI154940, NIH R01 EB010039, NIH R01 CA185251, NIH R01 CA186134, NIH R01 CA181648, NIH R01AI132627, NIH P30CA014520, EPA H-MAP 83573701, and American Cancer Society IRG-15-213-51.

We thank Mark McBride’s lab for providing strains and plasmids for this work. We also thank the DNA Sequencing Facility at University of Wisconsin-Madison Biotechnology Center for their next generation sequencing services. We gratefully acknowledge UW-Madison CALS Statistical Consulting for their assistance and thank Dr. Mark McBride and Dr. Edward Ruby for their helpful reviews of a previous version of the manuscript.

## AUTHOR CONTRIBUTION

S.M and J.H. designed the research project; S.M performed most of the experiments and A.H, J.F.N, M.G.C, C.L, and J.S contributed experimental expertise; S.M, M.G.C, and J.S analyzed data; J.H and D.J.B. acquired funding; S.M, A.H, J.F.N, M.G.C, C.L; and J.S provided methodology; S.M and J.H. prepared the original draft of the manuscript; S.M, A.H, J.F.N, M.G.C, C.L, J.S, and D.J.B reviewed and edited the manuscript.

